# An improved catalogue for whole-genome sequencing prediction of bedaquiline resistance in *M. tuberculosis* using a reproducible algorithmic approach

**DOI:** 10.1101/2025.01.30.635633

**Authors:** Dylan Adlard, Lavania Joseph, Hermione Webster, Ailva O’Reilly, Jeffrey Knaggs, Tim EA Peto, Derrick W Crook, Shaheed V Omar, Philip W Fowler

## Abstract

Bedaquiline (BDQ) has only been approved for use for a little over a decade yet is a key drug for treating multi-drug resistant tuberculosis, however rising levels of resistance threaten to reduce its effectiveness. Catalogues of mutations associated with resistance to bedaquiline are key to detecting resistance genetically for either diagnosis or surveillance. At present building catalogues requires considerable expert knowledge, often requires the use of complex grading rules, and is an irreproducible process. We developed an automated method, catomatic, that associates genetic variants with resistance (or susceptibility) using a two-tailed binomial test with a stated background rate and applied it to a dataset of 11,867 *Mycobacterium tuberculosis* samples with whole genome and bedaquline susceptibility testing data. Using this framework we investigated how to best classify variants and the phenotypic significance of minor alleles. The genes *mmpS5* and *mmpL5* are not directly associated with bedaquline resistance, and our catalogue of *Rv0678, atpE*, and *pepQ* variants attains a cross-validated sensitivity and specificity of 79.4 ± 1.8 % and 98.5 ± 0.3%, respectively, for 94 ± 0.4% of samples. Identifying samples with subpopulations containing *Rv0678* variants improves sensitivity, and detection thresholds in bioinformatic pipelines should therefore be lowered. By using a more permissive and deterministic algorithm trained on a sufficient number of resistant samples we have reproducibly constructed an AMR catalogue for BDQ resistance-associated variants that is comprehensive and accurate.

**Impact Statement:** Bedaquiline has recently received global endorsement for tuberculosis treatment, yet the genetic determinants of antimicrobial resistance remain incompletely understood. Existing gold-standard methods for building mutation catalogs lack public accessibility and reproducibility. We introduce catomatic, a reproducible and publicly available method that employs simpler statistics to increase sensitivity to resistance-associated variants. This approach has enabled investigations into mechanisms of resistance, the significance of genetic subpopulations, and key data attributes that influence the ease of classifying effects and the accuracy of BDQ resistance phenotype prediction in clinical samples. We strongly emphasise the utility in using reproducible statistics and sustainably developed software in genetics-focussed microbiology.

## Introduction

Bedaquiline (BDQ) is a diarylquinoline antimycobacterial agent that was approved for treatment of multidrug resistant tuberculosis (MDR-TB) by the US FDA in 2012. South Africa has a high burden of MDR-TB and rapidly pioneered the use of BDQ to treat MDR-TB patients, starting in late 2012^1^. Five years later in 2017 South Africa adopted a WHO-approved nine-month BDQ-containing regimen to treat patients with MDR-TB with no previous exposure to second-line treatments and who did not have extensive pulmonary or severe extra-pulmonary TB^2^. Bedaquiline contributed to the replacement of kanamycin, as part of a shift toward all-oral regimens – kanamycin is not orally bioavailable and is linked to numerous adverse effects^3–5^. More recently the WHO has endorsed a new six-month regimen for MDR-TB comprising bedaquline, pretomanid, linezolid, and moxifloxacin (BPaLM)^2^, and around 40 countries, including South Africa, have to date embraced this new regime^6^.

Patients with rifampicin-resistant TB (RR-TB) or MDR-TB who are treated with regimes containing BDQ can expect better outcomes; these include higher absolute rates of treatment success, a lower risk of mortality post treatment, higher sputum culture conversion rates, and reduced loss to follow-up^7–13^. Despite BDQ’s comparative success, antibiotic susceptibility testing (AST) is often not available and empirical treatment remains common^14^. Surveillance studies in South Africa estimate population or baseline levels of BDQ-resistance at 3.5-5.0%^1,15^, while treatment emergent resistance rates are variably reported between 2.0% and 6.0%^1,9,15^. The prevalence of resistance will no doubt have increased since these studies were published and a recent troubling correlation with rifampicin resistance has also been observed in Mozambique^16^.

Bedaquiline inhibits the membrane-embedded rotor of the mycobacterial ATP synthase (Fig. 1), encoded by the *atpE* gene, and therefore interfers with the production of ATP leading to cell death^17^. The WHO has identified *atpE*, together with *mmpL5, mmpS5, Rv0678*, and *pepQ*, as genes associated with resistance to BDQ (so-called Tier-1 genes), and *Rv1979c* as a gene possibly associated with resistance (a Tier-2 gene)^18^. Presumably because of the high likelihood of mutations in *atpE* introducing an unacceptably high fitness cost for the bacterium, few mutations in *atpE* have been observed; clinical and *in vitro* data instead suggest that resistance is most likely to arise through mutations in *Rv0678*, the MmpL5-MmpS5 efflux pump repressor^1,15,19–21^. For example a recent bacterial evolution study found that 86% of genetic variants that spontaneously arose under selection pressure due to the presence of BDQ were located in *Rv0678*, with only 8% of mutations arising in *atpE*^14^. The MmpL5-MmpS5 efflux pump is thought to export a range of compounds, including bedaquiline and clofazimine^22^, therefore it is unsurprising that mutations which abrogate the action of its repressor, *Rv0678*, compromise BDQ effectiveness. As yet no other genes have been identified by genome-wide association studies^23,24^

**Figure 1:**
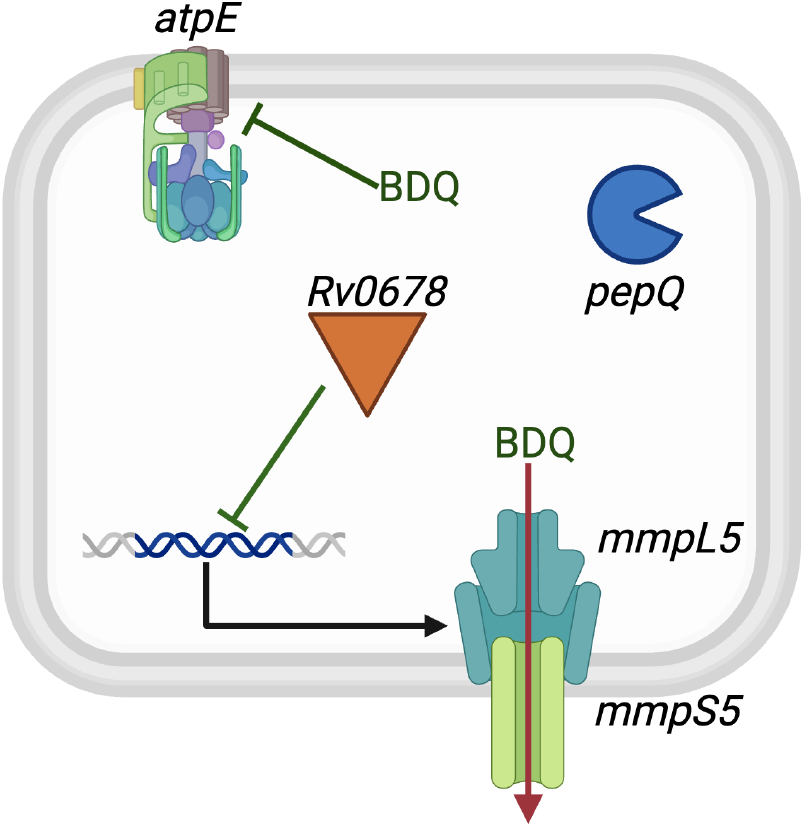
The genes considered to be associated with resistance to bedaquiline in *M. tuberculosis*. Bedaquline binds to the stalk of the ATPase which is encoded by the *atpE* gene. Resistance can arise when the drug is exported out of the cell by the MmpL5/MmpS5 efflux pump whose expression is regulated by *Rv0678*. Whilst *pepQ* has been implicated in resistance, the mechanism remains unclear; it encodes an aminopeptidase potentially affecting susceptibility via indirect effects on bacterial metabolism. Mutations in *Rv1979c*, which encodes a putative drug transporter, have also been associated with resistance, perhaps through altered efflux of BDQ, but this is not shown.

Bedaquiline was not included in the first edition of the WHO catalogue of genetic variants associated with resistance^25^ because there were insufficient resistant samples in the training dataset. Since then the WHO has amassed a dataset of 14,135 clinical samples (Table 1) that have undergone both whole genome sequencing and bedaquline phenotype drug-susceptibility testing and 1,032 (7.3%) of these are resistant. This dataset permitted the WHO to include BDQ in the second edition of the WHO catalogue (WHOv2) which was released in late 2023^18^. A total of 19 genetic variants were statistically associated with resistance to BDQ, resulting in a reported sensitivity of 49.4%, a specificity of 98.7% and a positive predictive value (PPV) of 75.2%^18^. The WHO applied a series of conservative statistical tests to each individual mutation^18^, thereby providing confidence that each association is correct but likely leads to false negative errors which may reduce the overall performance of the catalogue. Finally, because the training dataset of samples is not yet publicly available and the catalogue is built using a manual process with subsequent input from an expert committee (who e.g. add additional grading rules) it is slow to make and not reproducible.

**Table 1:** The number of samples in the Entire dataset, including the proportion of resistance and the phenotypic methods available. The dataset was created by aggregating samples from the CRyPTIC project with a smaller number collected by the NICD in South Africa. All CRyPTIC samples had BDQ minimum inhibitory concentrations measured using one of two bespoke 96-well broth microdilution plates (BMD)^33^.

| Source | Total samples | Resistant samples | %Resistant | Phenotypic methods |
| --- | --- | --- | --- | --- |
| CRyPTIC<sup>27</sup> | 4,387 | 29 | 0.7 % | UKMYC5 BMD plate |
| CRyPTIC<sup>27</sup> | 6,288 | 54 | 0.9 % | UKMYC6 BMD plate |
| NICD | 1,163 | 783 | 67.3% | MGIT960 |
| Total | 11,867 | 866 | 7.3% | BMD & MGIT960 |

Whole genome sequencing (WGS) infers the consensus genome from many smaller genetic fragments (reads), each of which may contain errors. To screen out these errors, bioinformatic pipelines usually only infer a mutation if it is supported by a majority of reads that map to the genetic locus under consideration^26^. For example the variant caller, clockwork, used by the WHO for the first edition of its catalogue^25^ and the CRyPTIC project^27^, by default requires 90% of reads for a genetic variant to be identified. This conservative approach implicitly assumes that the sample bacterial population is genetically homogeneous (which is not always true) and it has been shown that allowing far fewer reads to identify resistance alleles can improve the sensitivity of predicting fluoroquinolone resistance whilst not significantly reducing the specificity^28^. Consequently the WHO not only reduced this threshold to 75% when building the second edition of their catalogue but also re-evaluated the performance of the catalogue when only 25% of reads were required for a genetic variant to be called. Notably the sensitivities of several drugs all increased by more than two absolute percentage points^18^ suggesting that identifying minor alleles are important for predicting resistance to some drugs; these are, in descending order, bedaquline (+10.2%), moxifloxacin (+4.6%), levofloxacin (+4.4%), clofazimine (+4.3%), capromycin (+3.1%), amikaicin (+2.6%), kanamycin (+2.3%) and pyrazinamide (+2%).

In 2015 *Walker et al*.^29^ implemented the definite defectives algorithm^30^ to build a catalogue of resistance-associated variants for *M. tuberculosis*, focussing on the four first-line antibiotics, the fluoroquinolones and the aminoglycosides. Their approach defined susceptible variants as those with *consistently* susceptible phenotypes which assumes a high penetrance of resistant variants, and a low probability of resistance for as yet unobserved mutations. This prior probability was effective because for most antituberculars resistance-associated variants (RAVs) occur in essential genes, usually leading to a corresponding manyfold increase in the minimum inhibitory concentration (MIC). In their dataset most compounds behaved like this; a notable exception was pyrazinamide where most RAVs occur in *pncA* which is a non-essential gene. The first two versions of the WHO catalogue make the same implicit assumption for all drugs when they assume “Group 3 (Uncategorized)” variants are susceptible when calculating the sensitivities and specificities. Drugs where RAVs are found in non-essential genes such as *pncA* or *embB*, however, typically perform less well under this framework as there is typically more genetic variability and thus more data are required to achieve confidence in each susceptible classification^16,31^.

In this paper we shall describe our algorithm, catomatic, and use it to build an accurate catalogue of genetic variants associated with bedaquline susceptibility and resistance. Since the entire process is handled by software it is fast, deterministic and reproducible and we shall deliberately use less conservative statistics at the level of each individual mutation with the aim of improving the overall performance of the catalogue. This will necessarily force us to consider whether all Tier-1 genes (*mmpL5, mmpS5, Rv0678, atpE, pepQ*) play a role in resistance to bedaquiline. Being able to rapidly build a catalogue will also permit us to consider the impact on performance of altering the proportion of reads at a genetic loci required to support a variant call. This work is enabled by a large dataset of clinical samples from South Africa biased towards BDQ resistance which we use to enhance and enrich the published CRyPTIC datatset which, due to when it was collected, has a low incidence of BDQ resistance. We emphasise that how AMR catalogues are built and made available is important, and we will demonstrate there is real utility in using a deterministic, simpler, and more permissive approach that can be readily reproduced by other researchers.

## Materials and Methods

The dataset for this study was constructed by pooling two separate collections of *M. tuberculosis* samples. The first is based on 21,057 isolates collected by the CRyPTIC project, each of which had a bedaquiline minimum inhibitory concentration (MIC) measured using a bespoke 96-well broth microdilution plate^32,33^. Since reading the growth of *M. tuberculosis* in broth microdilution 96-well plates is a difficult and subjective task, we only took forward MICs where two or more independent methods^34,35^ agreed on the value to minimise measurement error^27,33^; this reduces the number of valid MICs to 14,605. Of these, 10,704 samples (Table 1) also underwent short-read (Illumina) whole genome sequencing (WGS) and only 83 samples (0.8 %) were resistant. This was determined by whether the MIC lies above a research epidemiological cut-off value (ECOFF) of 0.25 mg/L derived using interval regression on a dataset of 11,838 samples for BDQ^33^. Two plate designs (UKMYC5 and UKMYC6) were used, each incorporating different but overlapping BDQ concentration ranges in a doubling dilution series across eight wells^33^. The UKMYC5 plates covered a range of 0.015–2.0 mg/L, while the UKMYC6 plates spanned 0.008–1.0 mg/L (an example of the latter is shown in Fig 5B).

The second set of samples was collected by the NICD (The National Institute For Communicable Diseases) in South Africa and comprised 1,163 samples, all of which underwent short-read WGS and are deposited in the European Nucleotide Archive with study accessions PRJEB55007 and PRJEB76547. All samples were tested for bedaquiline resistance using the BACTEC MGIT960 system with the WHO-endorsed ECOFF of 1 mg/L. A subset (648 samples) also had their BDQ MIC measured using the BACTEC MGIT960 system. Overall 783 samples were assessed as being resistant by at least one method. The MICs in the NICD dataset are biased since there was strong sampling pressure for those assessed as being resistant to BDQ. Together, our dataset therefore comprises 11,867 samples of which 866 (7.3%) were assessed as resistant to BDQ – we call this the Entire Dataset.

The consensus genome for each of the 11,867 samples was inferred from the genetic short reads using the minos variant caller as incorporated into v0.12.4 of the clockwork pipeline^33,36^. The resulting variant call files (VCF) were then processed by gnomonicus (v2.5.1)^37^ which calculated all the resulting genetic variants in all Tier-1 genes; *mmpL5, mmpS5, pepQ, Rv0678* and *atpE*. An option was set to ignore the filter in the VCF files that prevent any variants supported by fewer than 90% reads being called. These minor alleles are annotated in the resulting data tables and the fraction of read support (FRS) also recorded. Wild-type samples, insertions or deletions (indels), synonymous and non-synonymous single nucleotide polymorphisms (SNPs), and premature stop codons were all identified on the basis of 90% or more of the reads supporting the genetic variant.

The Entire Dataset set of 11,867 genomes contains 25,651 mutations in the relevant genes (Table 2). The majority of these (21,416, 83.5 %) are phylogenetic mutations, such as T794I, D767N, and I948V in *mmpL5*^38^ and a small group (1,186) are synonymous. Because phylogenetic mutations have no impact on DST results,both are assumed to have no effect on the action of bedaquline and, since their inclusion would constrain the definite defectives algorithm by adding considerable noise, we created a smaller dataset of 2,116 samples all of which have at least one non-synonymous substitution, insertion or deletion in the Tier-1 genes. We call this the Training Subset; note that due to having too few samples resistant to bedaquiline, we do not have an independent Validation dataset. Almost half (1,455, 47.7 %) of the mutations in the Training dataset are in *Rv0678*, with 32.7% (983) in *mmpL5*.

**Table 2:** The number of samples with mutations in any of the WHO-defined BDQ candidate genes, and the number of mutations in the training and validation sets. The training set exclusively constitutes samples that contain non-synonymous, non-phylogenetic mutations, and all WT samples were filtered out. The validation set contains all mutations observed, as well as samples that contain no relevant mutations.

|  | Entire Dataset |  |  |  |  |  | Training Subset |  |  |  |  |  |
| --- | --- | --- | --- | --- | --- | --- | --- | --- | --- | --- | --- | --- |
|  | Samples |  |  | Mutations |  |  | Samples |  |  | Mutations |  |  |
| Gene | R | S | Total | R | S | Total | R | S | Total | R | S | Total |
| *Rv0678* | 709 | 451 | 1160 | 967 | 522 | 1489 | 709 | 426 | 1135 | 961 | 494 | 1455 |
| *pepQ* | 25 | 278 | 303 | 31 | 709 | 740 | 25 | 200 | 225 | 31 | 414 | 445 |
| *mmpL5* | 860 | 10902 | 11762 | 1637 | 21413 | 23050 | 9 | 784 | 793 | 17 | 966 | 983 |
| *mmpS5* | 4 | 80 | 84 | 5 | 84 | 89 | 3 | 57 | 60 | 4 | 60 | 64 |
| *atpE* | 15 | 30 | 45 | 15 | 169 | 184 | 14 | 17 | 31 | 14 | 88 | 102 |
| None | 5 | 94 | 99 | 5 | 94 | 99 |  |  |  |  |  |  |
| Total | 866 | 11001 | 11867 | 2660 | 22991 | 25651 | 733 | 1383 | 2116 | 1027 | 2022 | 3049 |

Building on the methodologies employed by Walker *et al*.^29^, WHOv1^25^, and WHOv2^18^, we applied the definite defectives algorithm^30^ to identify and classify benign variants. Our method differs by using a single, two-tailed binomial test to classify resistant and benign variants under the null hypothesis that there is no statistically significant difference between the proportion of resistance in samples containing the mutation, when it is the only mutation present across all candidate genes, and a specified background rate (Fig. S1A). We arbitrarily chose a background rate of 10%: this is conservative since it is greater than both the naive rate of 1.4% calculated from the dataset with no non-synonymous mutations in any of the candidate genes and reasonable estimates of measurement and labeling error.

If there is a predominance of susceptible samples at 95% confidence (inferred via a Wilson score interval), that particular mutation is classified as benign (susceptible) and removed from the Training Subset, potentially revealing additional samples with a single remaining mutations (Fig. S1A). This relies on the assumption that mutations that do not cause resistance can co-occur with those that do, and if a mutation in isolation does not cause resistance, then it will also not contribute to the phenotype when not in isolation.

At the point that all classifiable susceptible variants have been catalogued, the remaining variants are tested to see if they can be classified as Resistant (R) using the same hypothesis test; all remaining mutations are classified as Unknown (U). We use a ternary classification system (R vs S vs U) to report low confidence mutations and/or those whose calculated proportions do not differ sufficiently from the background.

Catalogues were saved as a CSV file in a format that can be parsed by a freely-available Python package, piezo,^39^, which was subsequently employed to make phenotypic predictions for each sample. Isolates containing at least one mutation catalogued as resistant were predicted to be resistant, while isolates containing no mutations, or only mutations catalogued as susceptible, were predicted to be susceptible. To demonstrate the utility of persisting ‘U’ mutation classifications through to sample phenotypic predictions, we employed two prediction protocols. The first method mirrors the WHO approach^18,25^ by assuming that samples containing uncatalogued and ‘U’ mutations are susceptible. The second protocol instead uses a ternary classification system (R vs U vs S) to report samples containing unseen and ‘U’ mutations (Fig. S1B). In both cases the sensitivity and specificity were calculated from a confusion matrix, however for the ternary system we additionally defined ‘definitive prediction rate (DPR)’ as the fraction of samples for which we can make a definite (i.e. R or S) prediction. Where cross-fold validation was carried out, scikit-learn^40^ was used to automate shuffled 5-fold cross-validation. Chi-squared (*χ*^2^) and Fisher’s exact tests are reported as appropriate..

### Data Summary

The catalogue construction algorithm was implemented using an open-source Python3 package we developed, catomatic,^41^ to allow for completely algorithmic and reproducible generation of catalogues. All data, results and most figures in this paper can be reproduced via a public GitHub repository (https://github.com/fowlerlab/tb-bdq-cat)^42^ that contains all necessary data and code, in the form of annotated Jupyter Notebooks.

## Results

### *mmpL5* and *mmpS5* are not resistance genes

Applying our algorithm to iteratively classify mutations in the Tier-1 candidate genes in the Training Subset assuming only 10% or more of reads are required to identify a genetic variant yielded 70 RAVs, seven susceptibility-associated variants (SAVs), and 459 genetic variants that could not be associated with either resistance or susceptibility (Fig. 2A). Of the 70 catalogued RAVs, 64 (91.4%) are in *Rv0678*, four (5.7%) are in *pepQ* and two (2.9%) are in *atpE* (Fig. 2A). The lack of any variants associated with a definite phenotype in the *mmpL5* and *mmpS5* genes mirrors other catalogues^18,25^ and does not justify their retention as their presence may mask the presence of other genetic variants. Removing both genes leads to a single additional RAV being identified in *Rv0678* (Fig. 2B). Accordingly, all catalogues constructed henceforth exclude *mmpL5* and *mmpL5* and therefore only *Rv0678, atpE* and *pepQ* are considered to be resistance genes.

**Figure 2:**
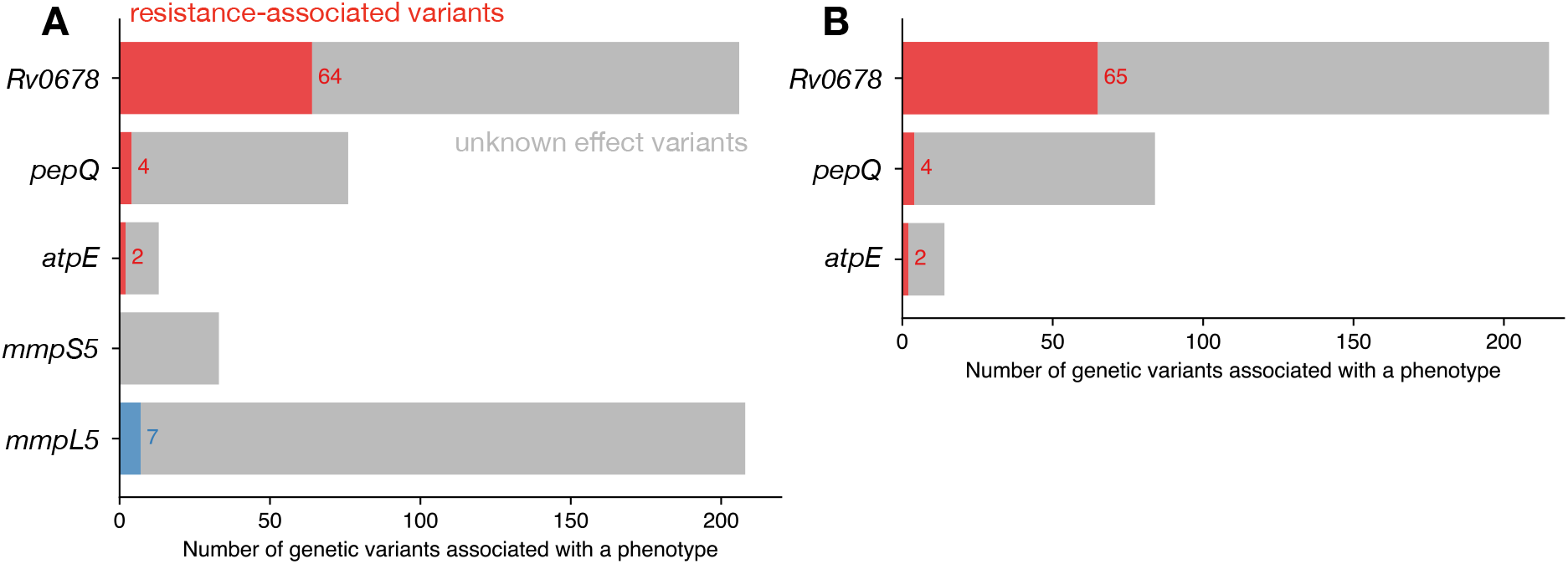
The majority of resistance associated variants in our initial catomatic catalogue occur in *Rv0678*. **A**. The number of resistant (R, red), susceptible (S, blue) and unclassified (U, grey) mutations in the catalogue trained on the Training subset when all Tier-1 candidate genes are included. No RAVs are found in *mmpS5* or *mmpL5*. **B**. Not considering either *mmpS5* or *mmpL5* leads to one additional RAV in *Rv0678* being identified.

### Resistance-associated variants are found throughout the entire Rv0678 protein

RAVs are found along the entire length of the *Rv0678* gene with no distinct localisation (Fig. 3A). Plotting them onto the structure of the Rv0678 protein (Fig. 3B) shows that several high-prevalence RAVs are identified within the DNA binding region (codons 34-99), accounting for 79.6% of catalogued RAVs, 49.8% of which were detected in the *α*2*α*3 helix. The majority of variants in the DNA binding region (62.8%) were insertions leading to a frame shift at codons 46, 47, and 64. Among the remaining 112 samples, 94 (17.1%) isolates harboured mutations in the dimerisation domain (codons 16-33 and 100-160), with the remaining 18 sitting outside known functional domains. Of the 71 rows in this initial catomatic catalogue, 26 (36.7%) describe frameshifts arising from insertions or deletions and five (7.04%) are premature stop codons; these account for 275 (41.6%) and 14 (2.1%) of the resistant samples in the Training Subset, respectively.

**Figure 3:**
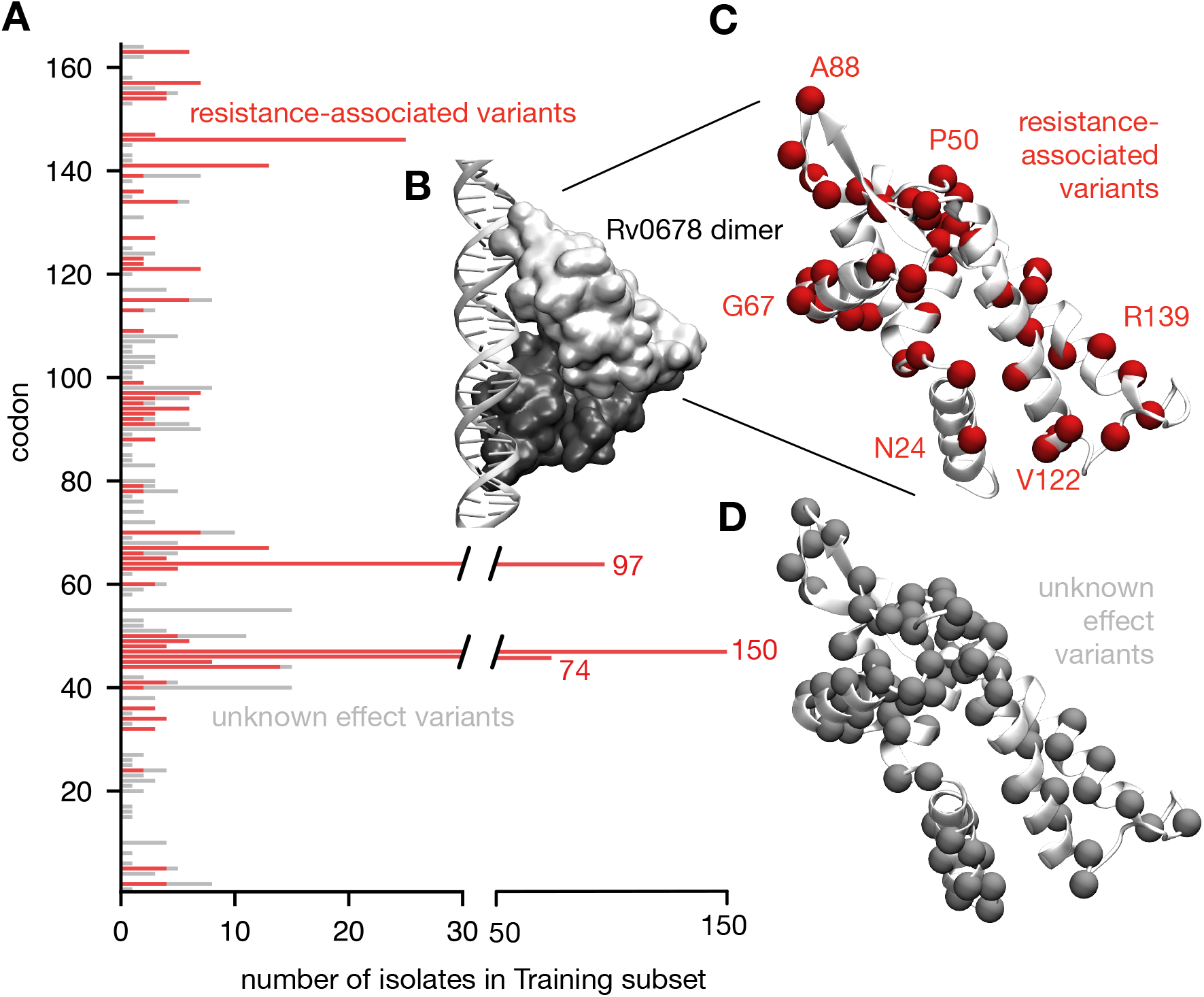
Resistance associated variants are found throughout the *Rv0678* gene. **A**. Whilst mutations at several codons in the *Rv0678* gene are associated with resistance in large numbers of isolates in the Training Subset (notably codons 46, 47 and 64), resistance to bedaquiline can arise from non-synonymous mutations along the entire gene. The number of isolates containing genetic variants that could not be associated with either resistance or susceptibility are drawn in grey. No genetic variants were associated with susceptibility. **B**. Rv0678 is a transcriptional regulator and the protein forms a dimer (monomers coloured light and dark grey). Shown in this figure is a model based on the experimental structure of how it binds to DNA^14^. **C**. Annotating the codons where variants have been associated with resistance suggests that there is possibly some correlation with structural features e.g. the *α*2*α*3 helix (containing Gly67) which intercalates into the major groove of the DNA in the model, but this is not conclusive. **D**. There is no clear difference when compared to which codons have variants not associated with either resistance or susceptibility (in grey).

### Our initial catalogue performs similarly to the second edition of the WHO catalogue

To assess the accuracy of our initial catomatic catalogue, we applied it to the Entire Dataset, which is a superset of the Training Subset and contains wild-type samples and samples containing synonymous and phylogenetic mutations (Table 2). By adopting the binary paradigm used by the WHO, whereby samples containing one or more catalogued R mutations are classified as resistant and all other samples as susceptible, we achieved a sensitivity of 70.1% (a false negative rate, or in clinical terms a very major error, VME, of 29.9%) and a specificity of 98.4% (a false positive rate, or a major error, ME, of 1.6%) (Fig. 4A, `catomatic-1`, Table 3). Applying WHOv2 to our Entire Dataset with the same minimum 10% threshold for calling variants yielded a sensitivity of 68.0% and specificity of 98.2% (Fig. 4E, WHOv2-1, Table 3), values which are encouragingly similar to `catomatic-1`, as evidenced by *χ*^2^ p-values of 0.38 and 0.14, respectively. Notably, the WHOv2 performance on our Entire Dataset is also similar to its original reported performance as calculated by applying it to its training dataset^18^.

**Table 3:** The performance of catomatic catalogues generated in this study. All catalogues (including WHOv2) were validated on the Entire Dataset. Catalogues were built and evaluated at specified and sometimes different Fractions of Read Support (FRS), allowing e.g. a catalogue built on effectively genetic homogenous samples (0.9) to be applied to samples that can contain minor alleles (0.1). To achieve a binary classification all samples predicted to have an unknown phenotype were assumed to be susceptible (WHOv2-1 & `catomatic-2a`), whilst the ternary logic permits all three values. This latter approach requires an additional metric, definitive prediction rate (DPR), which is the proportion of samples for which resistant or susceptible predictions can be made. The Run number is for reference: `catomatic-2a` is our recommended catalogue and prediction logic, whilst `catomatic-2b` mirrors `catomatic-2a` but used five-fold cross-validation to estimate the uncertainty in the performance, and `catomatic-3` included generalisable, manually added loss of function rules that replace all other indel or premature stop rows.

| Catalogue | Run | Fraction of Read Support |  | Logical paradigm | R:S (#) | Sensitivity (%) | Specificity (%) | DPR (%) |
| --- | --- | --- | --- | --- | --- | --- | --- | --- |
|  |  | Build | Evaluated |  |  |  |  |  |
| WHOv2 | 1 | 0.75 | 0.10 | binary | 19:0 | 68.0 | 98.2 | - |
| *catomatic* | 1 | 0.1 | 0.10 | binary | 71:0 | 70.1 | 98.4 | - |
| WHOv2 | 2 | 0.75 | 0.10 | ternary | 19:0 | 80.4 | 98.1 | 96.0 |
| WHOv2 | 3 | 0.75 | 0.75 | ternary | 19:0 | 56.1 | 98.7 | 96.8 |
| WHOv2 | 4 | 0.75 | 0.25 | ternary | 19:0 | 76.9 | 98.3 | 96.1 |
| *catomatic* | 2a | 0.1 | 0.10 | ternary | 71:0 | 81.8 | 98.4 | 95.1 |
| *catomatic* | 2b | 0.1 | 0.10 | ternary | 51-60:0 | $79.4 \pm 1.8$ | $98.5 \pm 0.3$ | $94.2 \pm 0.4$ |
| *catomatic* | 3 | 0.1 | 0.10 | ternary | 36:0 | 83.2 | 98.1 | 95.9 |
| *catomatic* | 4 | 0.75 | 0.75 | ternary | 67:0 | 60.0 | 97.9 | 96.8 |
| *catomatic* | 5 | 0.75 | 0.25 | ternary | 67:0 | 78.3 | 97.6 | 95.9 |
| *catomatic* | 6 | 0.9 | 0.10 | ternary | 59:0 | 81.2 | 97.5 | 95.7 |

**Figure 4:**
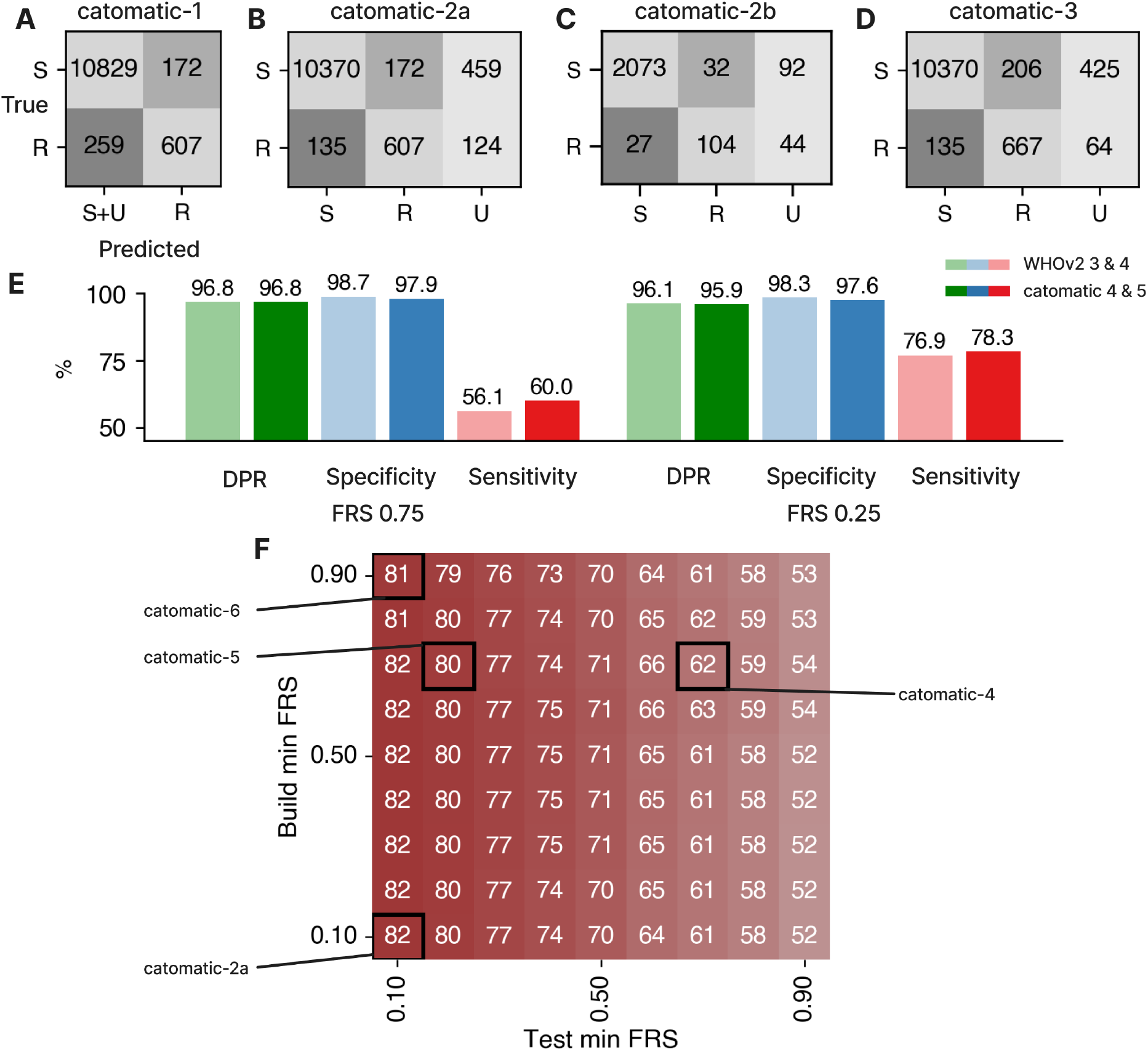
**A-D**. Confusion matrices for performance of the catalogue on the validation dataset, where the y axis represents actual phenotypes and the x axis represents predictions. **A**. Performance when a binary prediction system is used and samples containing U mutations are assumed susceptible (`catomatic-1`). **B**. Performance when a ternary prediction system is used (i.e. samples that hit only S and U rules are predicted ‘U’) (`catomatic-2a`). **C**. Performance when a ternary prediction system is used, under a five-fold cross validation strategy (`catomatic-2b`). **D**. Performance using a ternary prediction system for the catalogue with additional arbitrary LoF rules for *Rv0678* and *pepQ* (`catomatic-3`).**E**. Performance comparison between the WHOv2 catalogue and the catalogue constructed by catomatic at FRS 0.75, validated on our dataset, at FRS 0.75 (WHOv2-3 vs `catomatic-4`) and FRS 0.25 (WHOv2-4 vs `catomatic-5`), following the ternary prediction system. **F**. Performance of the catomatic catalogue when evaluated *and constructed* as a function of minimum FRS at 0.10 increments, using the ternary prediction logic. Black boxes represent the 0.10 increments in which `catomatic-2a`, `catomatic-4`, `catomatic-5` and `catomatic-6`’s minimum FRS exist. Specificity and DPR plots to be found in Fig. S7.

### Genetic variants not able to be associated with a definite phenotype should not be assumed to be susceptible to bedaquiline

This approach assumes that samples predicted to have an Unknown phenotype should be treated as Susceptible and we argue that this is inappropriate for bedaquiline where, despite best efforts, the sensitivity remains moderate, indicating that there are genes and/or genetic variants yet to be characterised in sufficient numbers that they can be associated with a definite phenotype, especially resistance. A ternary classification system is a natural consequence of using the two-tailed test and should hence be maintained through to the final prediction for each sample. We shall accordingly introduce an additional performance metric, definitive prediction rate (DPR): this is the fraction of samples for which definitive (R or S) predictions can be made. This approach results in a sensitivity of 81.8% (VME 18.2%) and a specificity of 98.4% (ME 1.6%) for 95.1% of the samples (Fig. 4B, `catomatic-2a` Table 3) and is a fairer reflection of the data. The ME and VME rates are attributable to 172 false positives and 135 false negatives, respectively. Clearly it is not best practice and could be mis-leading to validate on a dataset that is not independent to the training set so, since we lack sufficient data to create such a test set, we performed five-fold shuffled cross-validation to estimate the uncertainty in the sample predictions. This yielded a sensitivity of 79.4 ± 1.8% and specificity of 98.5 ± 0.3% for 94.2 ± 0.4% of samples (Fig. 4C, `catomatic-2b`, Table 3).

### Analysing the discrepants is complex and suggests there may be resistance mechanisms and genes not yet identified

The false positive predictions derive in part from samples that contain minor alleles containing indels, such as the single base insertions at codons 46, 47, and 64 in *Rv0678*. Of the 382 samples harboring a single catalogued mutation that could plausibly lead to loss of function (LoF), only 289 (75.7%) are phenotypically resistant, and notably only 43.2% of those containing an insertion at codon 64 are resistant (Fig. S2). It is not particularly instructive to analyse their respective MIC distributions because the majority of the samples measured to be resistant to BDQ occur in the NICD dataset which is effectively truncated since, by definition, it contains few samples with a low BDQ MIC since the samples were selected for resistance (Fig. S8).

Another complicating factor is that a linkage between LoF mutations in *mmpL5* and RAVs in *Rv0678* has been reported, whereby alleles with a dysfunctional pump are susceptible, irrespective of *Rv0678* variation^18,43^. Although not identifiable in the NICD dataset, we identify 54 such samples in the CRyPTIC data, and these primarily have a LoF mutation at codon 201 in *mmpL5*, 39 of which also contain an insertion at codon 64 in *Rv0678*. All these samples are susceptible to BDQ, with the phenotype linkage being statistically significant (Fisher’s exact p-value = 0.00034). This partially explains the heteroresistance observed at codon 64 and taking account of this effect increases the proportion of resistant samples harboring a single LoF mutation at this position from 43.2% to 84.6%.

The prevalence of putative LoF mutations in *Rv0678* prompted us to try adding a general rule associating any and all LoF mutations with resistance, similar to the approach used in WHOv2^18^. However, most LoF variants had already been catalogued, and so this led to only a slight, non-significant increase (+1.4%, *χ*^2^ p-value = 0.52) in sensitivity as 60 additional samples were correctly predicted resistant, and a slight decrease (−0.3%, *χ*^2^ p-value = 0.093) in specificity (Fig. 4D, `catomatic-3`, Table 3). We therefore have erred on the side of both caution and reproducibility and have refrained from including the rule in our final catomatic catalogue.

Among the 135 false negative samples, 131 contain no non-phylogenetic mutations in any of the Tier-1 candidate genes, and the remaining four samples contain non-synonymous, non-phylogenetic mutations in *mmpL5* (A736D, A755T, S721N, L709I). The majority (109) derive from the NICD dataset, suggesting either a resistance mechanism or gene is missing, there are labeling or phenotyping errors or both.

### Allowing minor alleles to contribute to classifying samples improves prediction performance

Genetic sub-populations (minor alleles), are typically filtered out in bioinformatic pipelines to minimize sequencing errors by insisting all genetic variants are supported by a specified proportion of reads at that locus (the Fraction of Read Support, FRS). The WHOv2 catalogue was constructed at a high FRS of 0.75 and then evaluated at the same FRS and also a lower value of 0.25. Allowing minor alleles to contribute led to a 10.2% increase in sensitivity on their training dataset^18^, a 14.5% increase on our Entire Dataset using their prediction logic, and a 20.8% increase on our data using the ternary system (WHOv2-3 & WHOv2-4, Table 3), indicating that our data contain more resistance-associated variants (RAVs) observed at both high *and* low FRS. We observed a similar effect for our catomatic catalogues on the Entire Dataset, with sensitivity increasing by 18.3% (*χ*^2^ p-value = 0.0) when the catalogue built at an FRS of 0.75 is evaluated at 0.25 (Fig. 4E, `catomatic-4` & `catomatic-5`, Table 3).

If minor variants are truly predictive, we should logically also *build* at a low fraction read support to capture those variants. Since our method is algorithmic, this is straightforward and we built and evaluated catalogues as a function of FRS (Fig. 4F). Decreasing the FRS we built at from 1 to 0.1 (fixing the evaluation FRS at 0.1) led to a small increase in sensitivity (0.6%, *χ*^2^ p-value = 0.82) owing to the classification of 14 additional RAVs, although two RAVs were also lost, one of which is c-11a, a high-frequency SNP in the *Rv0678* promoter observed in nine resistant samples at high FRS and 93 susceptible samples at lower FRS. However, this performance gain was minor, suggesting the read support used to build the catalogues makes little difference as most RAVs have already been captured at high FRS. However, because RAV’s do exist at low FRS, dropping the threshold when evaluating has a considerable effect on performance, and hence *the evaulation FRS is important* and should be lowered.

### Samples with a low fraction of reads supporting resistance have the same minimum inhibitory concentration and grow as well on the 96-well plate as homogenous samples

We hypothesised that resistant alleles when present in a fraction of a mixed sample would induce less resistance due to the fixed incubation time used in culture-based testing. We accordingly examined whether there is a correlation between FRS and MIC after two weeks incubation across all isolates with only one *Rv0678* resistant mutation (and any number of susceptible mutations) and specifically for *Rv0678* 141-ins-c, for which we have a reasonable number of samples. This analysis is only possible for the CRyPTIC samples which were grown on 96-well plates (Table 1) as both MIC and the % growth in all wells are available. Our results suggest that the proportion of reads describing any minor allele has no discernable effect on MIC (Fig. 5A, B) in the range 10-100%. We can also test if there is a relationship between FRS and bacterial growth. Plotting FRS versus percentage growth 14 days post-incubation, averaged over well concentrations of 0.015 and 0.03 mg/L (i.e., the two lowest values shared between the UKMYC5 and UKMYC6 plate designs) for all isolates with only one resistant mutation, and for *Rv0678* 141-ins-c, returned Pearson correlation coefficients of −0.04 and −0.08, respectively (Fig. 5B, C), suggesting that the proportion of a mixture containing an RAV has no impact on growth after two weeks incubation or the level of resistance against BDQ. Hence even samples containing small resistant subpopulations appear able to grow sufficiently fast enough to be indistinguishable from homogenous resistant samples after two weeks incubation.

**Figure 5:**
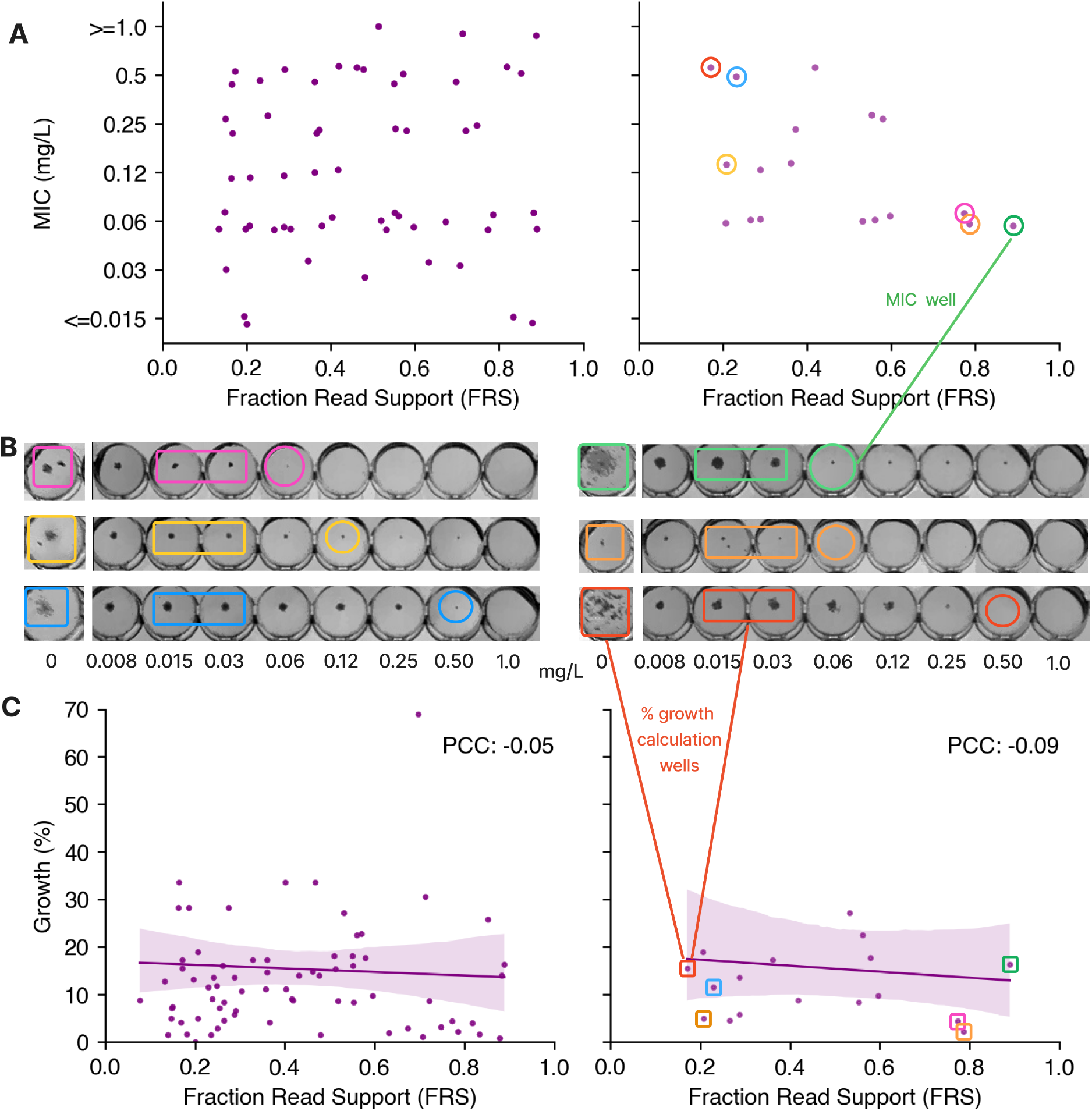
The fraction of read support (FRS) for a minor RAV does not correlate with either MIC or plate growth. **A**. FRS vs MIC (FRS < 0.9) for all UKMYC samples containing a single variant in *Rv0678* (first panel), and for all samples containing only 141-ins-c in *Rv0678* (second panel), with vertical jitter applied within each MIC interval for clarity. Note that UKMYC5 and UKMYC6 well concentrations have different ranges, and MICs have thus been aggregated into a single axis by truncating extreme MICs into the closest shared value, and therefore does not, for example, align with the MIC range in panel **B** which represents UKYMYC6 labels only. **B**. The corresponding images of 6 example UKMYC6 plated samples at the lower and upper ends of the FRS range. The first column is the control well from which percentage growth was calculated, while subsequent wells are increasing BDQ doubling dilutions. Circled are plotted MICs, and rectangles are growth values used to calculate percentage growth. **C**. FRS vs percentage growth on culture plates (FRS < 0.9) for all samples containing a single variant in *Rv0678* (first panel), and for all samples containing only 141-ins-c in *Rv0678* (second panel). Growth was calculated by averaging the lowest 2 well concentrations containing growth that are shared across plate designs, 0.015 and 0.03 mg/L.

## Discussion

We have developed a robust and reproducible method to iteratively classify the effect of mutations on the action of bedaquline and thereby build a comprehensive resistance catalogue. Our approach classified 65 resistance-associated variants (RAVs) in *Rv0678*, four in *pepQ*, and two in *atpE*, achieving a cross-validated performance of 79.4 ± 1.8% sensitivity and 98.5% ± 0.3% specificity, covering 94.2 ± 0.4% of samples (Table 3). Notably, our catalogue is the first to suggest it is at least possible to meet the 80% sensitivity threshold set by the WHO’s 2023 target product profiles for BDQ drug susceptibility testing^44^. Our method diverges from the WHO’s by using simpler, more inclusive statistical criteria when classifying individual genetic variants, helping enable the classification of 37 additional mutations. We also underscore the critical need for careful consideration of unclassified mutations.

RAVs are distributed along the entire length of the *Rv0678* gene, with a dominant cluster in the DNA-binding domain, reflecting the results of recent *in vitro* evolution models^14,45^ (Fig. 3). Despite being listed as candidate genes, we and others^1,14,15^ found no evidence directly linking *mmpL5* or *mmpS5* genetic variants to resistance, which is consistent with their biology; up-regulation of the efflux pump through disruption of its negative regulator (*Rv0678*) is a more plausible mechanism than somehow increasing the transport of BDQ via a point mutation. However, should the pump be disabled by a loss of function mutation in *mmpL5*, no variation in *Rv0678* can confer resistance as BDQ can no longer be exported. This co-occurrence renders *Rv0678* mutations functionally irrelevant^18,43^, and we observe the effect in 54 samples from the CRyPTIC dataset.

Bedaquline has only been used clinically for less than a decade and the prevalence of resistance remains, on average, low which has constrained the number of resistant samples in our dataset, limiting our ability to classify genetic variants. Additionally, the moderate sequencing depth typically present in our dataset will have prevented us resolving the presence of resistant minor alleles present at very low proportions. As a consequence, statistical power was insufficient to offset the test’s bias toward detecting resistance, restricting the catalogue to resistant classifications only. Dramatically expanding the size of our dataset would not only enable independent validation of catalogues but also facilitate classification of susceptible genetic variants, moving us closer to being able to predict when bedaquline should be administered, rather than solely when it should be avoided^29^.

Simply labelling each sample as resistant or susceptible in theory enables larger datasets to be created by aggregating smaller datasets that use different phenotypic drug susceptibility testing methods without needing to account for methodological differences. However, here we merged a larger dataset from a time period where there was minimal BDQ use with a smaller dataset of more-recently collected resistant samples (Fig. S9). This complicates calculating a meaningful background resistance rate, as the larger and smaller datasets are skewed towards susceptibility and resistance, respectively. Consequently, we arbitrarily set the background rate at 10% and encourage researchers to carefully choose the background rate depending on what they are seeking to optimise. For example, raising the background rate will permit additional susceptible mutations to be classified but increases the risk of predicting a resistant infection as susceptible (a so-called Very Major Error).

Merging datasets requires that the drug susceptibility methods give equivalent results. We do not have any samples tested on both a UKMYC plate and MGIT but we can compare samples (Fig. S8) with a single, specified mutation in *Rv0678* (and no non-synoymous mutations in any of the other bedaquiline resistance genes). Unfortunately since all the samples which had MICs measured using MGIT were enriched for BDQ resistance, this has the effect of biasing these MIC distributions making comparison with the UKMYC dataset difficult. That said, examining the two most common frameshifts (141 ins c and 138 ins g) suggests that, at the very least, the MICs are not equivalent and the ECOFF/ECVs may not be equivalent between the methods. This picture is not supported by the rarer mutations. Overall the picture is complex and we cannot be certain both methods are equivalent due to the small sample sizes and the lack of a direct comparison. Because the MGIT samples were all collected in one laboratory it is also possible there are other genetic effects (e.g. lineage or resistance to other drugs) affecting the MICs that we have not taken into account.

Another potential source of heterogeneity in the reported binary phenotypes is that genetic variation in *Rv0678* appears to only modestly elevate MIC, often leading to MIC distributions for specific genetic variants that overlap the breakpoint^14,45^. It has even been suggested that LoF mutations in *Rv0678*, which constitute the most frequent RAVs, can shift the gene into a reading frame that retains wild-type functionality^46^. While our UKMYC plate MIC distributions support this, the LoF-containing MGIT samples sit firmly above the ECOFF (Fig. S8). A limitation of purely relying a binary or ternary classification system is that it becomes difficult to distinguish a biological explanation for heterogeneity from phenotype artifacts.

The observed modest increases in MIC are likely a consequence of a limited fitness cost accompanying the variation, which is consistent with *Rv0678* being non-essential. The relative lack of a fitness cost implies a small subpopulation of bacteria with an RAV could quickly outcompete the majority wild-type population under bedaquiline treatment, which is consistent with the lack of any correlation between the fraction of genetic reads supporting a resistant *Rv0678* allele (Fig. 5) and the observed bacterial growth after two weeks incubation. When applying a resistance catalogue to detect bedaquiline resistance, it would therefore seem sensible to detect minor alleles containing RAVs, as recommended for the fluoroquinolones^28^. We note that we have not been able to set a minimum read threshold in this work and elucidating this will be important.

The absence of what one might call “classical” bimodality in the bedaquiline MIC distribution suggests that a regression approach that can take account of MIC being an interval may be more appropriate^16^. This would resolve the problems with binary heterogeneity and avoid having to assume an arbitrary background resistance rate. Since each MIC would have a confidence limit, applying a threshold to predicted MICs would naturally lead to, at the very least, three classification labels (Resistant/Unknown/Susceptible). Furthermore, regression with interaction terms would inherently account for linkage effects, such as those observed between LoF mutations in *mmpL5* and *Rv0678*.

The key strength of catomatic is speed; it can generate or update catalogues from tables of processed genetic and phenotypic data in minutes. When coupled with a robust WGS pipeline this enables the rapid creation of catalogues based on specific sample collections perhaps allowing current resistance patterns to be detected — including for novel drugs if phenotyping is available — or permitting catalogues tailored to specific geographies or lineages to be created. In future work will automatically build a catalogue for all a panel of TB drugs, including validating its performance on a hold-out dataset and comparing that to the performance of the WHO endorsed catalogues. Given its speed and reproducibility, our approach offers a promising alternative to the current process used by the WHO to build their resistance catalogues.

In summary, five key factors simplify the creation of a resistance catalogue: resistance genes being essential, mutations causing a significant increase in MIC, a high prevalence of resistance within the training dataset, concordance between DST methods when multiple are used, and having a large dataset to work with. Bedaquiline is a challenge as these factors are either not true or they are yet to be met. Regardless of the statistical approach, it is crucial that researchers rigorously validate and present their catalogues. We stress the need for fully reproducible statistics and sustainable software practices. Our approach is algorithmic, deterministic and requires minimal domain expertise and user intervention. We hope the discipline will move towards adopting reproducibility best practices *viz*. be held under version control together with all derivation code and the training and validation datasets. Ideally, the catalogue itself should be a single artefact that can be easily parsed by both humans and computers to enable further work. There are many barriers preventing the adoption of WGS in low-resource settings where MDR-TB is more likely to be prevalent, however adopting these standards will both help boost the global uptake of existing molecular diagnostics (including WGS) and the development of new methods.

## References

[1] Pai, H., Ndjeka, N., Mbuagbaw, L., Kaniga, K., Birmingham, E. et al. Bedaquiline safety, efficacy, utilization and emergence of resistance following treatment of multidrug-resistant tuberculosis patients in south africa: a retrospective cohort analysis. BMC Infectious Diseases 22 (2022).

[2] World Health Organisation. Rapid communication: Key changes to the treatment of drug-resistant tuberculosis (2022).

[3] Department of Health South Africa. Interim clinical guidance for the implementation of injectable-free regimens for rifampicin-resistant tuberculosis in adults, adolescents and children (2018).

[4] Falzon, D., Schünemann, H. J., Harausz, E., González-Angulo, L., Lienhardt, C. et al. World health organization treatment guidelines for drug-resistant tuberculosis, 2016 update. European Respiratory Journal 49, 1602308 (2017).

[5] World Health Organisation. WHO treatment guidelines for drug-resistant tuberculosis, October 2016 revision.

[6] World Health Organisation. Global tuberculosis report 2023 (2023).

[7] Wang, M.-G., Wu, S.-Q. & He, J.-Q. Efficacy of bedaquiline in the treatment of drug-resistant tuberculosis: a systematic review and meta-analysis. BMC Infectious Diseases 21, 970 (2021).

[8] Ndjeka, N., Campbell, J. R., Meintjes, G., Maartens, G., Schaaf, H. S. et al. Treatment outcomes 24 months after initiating short, all-oral bedaquiline-containing or injectable-containing rifampicin-resistant tuberculosis treatment regimens in south africa: a retrospective cohort study. The Lancet Infectious Diseases 22, 1042–1051 (2022).

[9] Nimmo, C., Millard, J., Brien, K., Moodley, S., van Dorp, L. et al. Bedaquiline resistance in drug-resistant tuberculosis hiv co-infected patients. European Respiratory Journal 55, 1902383 (2020).

[10] Zhao, Y., Fox, T., Manning, K., Stewart, A., Tiffin, N. et al. Improved treatment outcomes with bedaquiline when substituted for second-line injectable agents in multidrug-resistant tuberculosis: A retrospective cohort study. Clinical Infectious Diseases 68, 1522–1529 (2019).

[11] Ahmad, N., Ahuja, S. D., Akkerman, O. W., Alffenaar, J.-W. C., Anderson, L. F. et al. Treatment correlates of successful outcomes in pulmonary multidrug-resistant tuberculosis: an individual patient data meta-analysis. The Lancet 392, 821–834 (2018).

[12] Olayanju, O., Limberis, J., Esmail, A., Oelofse, S., Gina, P. et al. Long-term bedaquiline-related treatment outcomes in patients with extensively drug-resistant tuberculosis from south africa. European Respiratory Journal 51, 1800544 (2018).

[13] Schnippel, K., Ndjeka, N., Maartens, G., Meintjes, G., Master, I. et al. Effect of bedaquiline on motality in south african patients with drug-resistant tuberculosis: a retrospective cohort study. The Lancet Respiratory Medicine 6, 699–706 (2018).

[14] Sonnenkalb, L., Carter, J. J., Spitaleri, A., Iqbal, Z., Hunt, M. et al. Bedaquiline and clofazimine resistance in Mycobacterium tuberculosis: an in-vitro and in-silico data analysis. The Lancet. Microbe (2023).

[15] Ismail, N. A., Omar, S. V., Moultrie, H., Bhyat, Z., Conradie, F. et al. Assessment of epidemiological and genetic characteristics and clinical outcomes of resistance to bedaquiline in patients treated for rifampicin-resistant tuberculosis: a cross-sectional and longitudinal study. The Lancet Infectious Diseases 22, 496–506 (2022).

[16] The CRyPTIC Consortium. Quantitative measurement of antibiotic resistance in Mycobacterium tuberculosis reveals genetic determinants of resistance and susceptibility in a target gene approach. Nature Communications 15 (2024).

[17] Preiss, L., Langer, J. D., Yildiz, O., Eckhardt-Strelau, L., Guillemont, J. E. G. et al. Structure of the mycobacterial ATP synthase Fo rotor ring in complex with the anti-TB drug bedaquiline. Science Advances 1, 1–9 (2015).

[18] World Health Organization. Catalogue of mutations in Mycobacterium tuberculosis complex and their association with drug resistance. Second edition (2023).

[19] Hartkoorn, R. C., Uplekar, S. & Cole, S. T. Cross-resistance between clofazimine and bedaquiline through upregulation of mmpl5 in Mycobacterium tuberculosis. Antimicrobial Agents and Chemotherapy 58, 2979–2981 (2014).

[20] Ismail, N., Ismail, N. A., Omar, S. V. & Peters, R. P. H. In vitro study of stepwise acquisition of rv0678 and atpe mutations conferring bedaquiline resistance. Antimicrobial Agents and Chemotherapy 63, 10.1128/aac.00292–19 (2019).

[21] Kadura, S., King, N., Nakhoul, M., Zhu, H., Theron, G. et al. Systematic review of mutations associated with resistance to the new and repurposed Mycobacterium tuberculosis drugs bedaquiline, clofazimine, linezolid, delamanid and pretomanid. Journal of Antimicrobial Chemotherapy 75, 2031–2043 (2020).

[22] Briffotaux, J., Huang, W., Wang, X. & Gicquel, B. MmpS5/MmpL5 as an efflux pump in Mycobacterium species. Tuberculosis 107, 13–19 (2017).

[23] Consortium, T. C. Genome-wide association studies of global Mycobacterium tuberculosis resistance to 13 antimicrobials in 10,228 genomes identify new resistance mechanisms. PLOS Biology 20, e3001755 (2022). Publisher: Public Library of Science.

[24] Coll, F., Phelan, J., Hill-Cawthorne, G. A., Nair, M. B., Mallard, K. et al. Genome-wide analysis of multi- and extensively drug-resistant Mycobacterium tuberculosis. Nature Genetics 50, 307–316 (2018).

[25] World Health Organisation. Catalogue of mutations in Mycobacterium tuberculosis complex and their association with drug resistance (2021).

[26] Mohammed, K. S., Kibinge, N., Prins, P., Agoti, C. N., Cotten, M. et al. Evaluating the performance of tools used to call minority variants from whole genome short-read data. Wellcome Open Research 3, 21 (2018).

[27] Crook, D. W., Peto, T. E., Hoosdally, S. J., Cruz, A. L. G., Walker, A. S. et al. A data compendium associating the genomes of 12,289 Mycobacterium tuberculosis isolates with quantitative resistance phenotypes to 13 antibiotics. PLoS biology 20, e3001721 (2022).

[28] Brankin, A. E. & Fowler, P. W. Inclusion of minor alleles improves catalogue-based prediction of fluoroquinolone resistance in Mycobacterium tuberculosis. JAC-Antimicrobial Resistance 5 (2023).

[29] Walker, T. M., Kohl, T. A., Omar, S. V., Hedge, J., Elias, C. D. O. et al. Whole-genome sequencing for prediction of Mycobacterium tuberculosis drug susceptibility and resistance: A retrospective cohort study. The Lancet Infectious Diseases 15, 1193–1202 (2015).

[30] Dorfman, R. The detection of defective members of large populations. The Annals of Mathematical Statistics 14, 436–440 (1943).

[31] Carter, J. J., Walker, T. M., Walker, A. S., Whitfield, M. G., Morlock, G. P. et al. Prediction of pyrazinamide resistance in Mycobacterium tuberculosis using structure-based machine-learning approaches. JAC-Antimicrobial Resistance 6, dlae037 (2024).

[32] Rancoita, P. M. V., Cugnata, F., Gibertoni Cruz, A. L., Borroni, E., Hoosdally, S. J. et al. Validating a 14-Drug Microtiter Plate Containing Bedaquiline and Delamanid for Large-Scale Research Susceptibility Testing of Mycobacterium tuberculosis. Antimicrobial Agents and Chemotherapy 62, e00344–18 (2018).

[33] The CRyPTIC Consortium. Epidemiological cutoff values for a 96-well broth microdilution plate for high-throughput research antibiotic susceptibility testing of M. tuberculosis. Eur Respir J 60, 2200239 (2022).

[34] Fowler, P. W., Gibertoni Cruz, A. L., Hoosdally, S. J., Jarrett, L., Borroni, E. et al. Automated detection of bacterial growth on 96-well plates for high-throughput drug susceptibility testing of Mycobacterium tuberculosis. Microbiology 164, 1522–1530 (2018).

[35] Fowler, P. W., Wright, C., Spiers-bowers, H., Zhu, T., Baeten, E. M. L. et al. A crowd of BashTheBug volunteers reproducibly and accurately measure the minimum inhibitory concentrations of 13 antitubercular drugs from photographs of 96-well broth microdilution plates. eLife (2022).

[36] Hunt, M., Letcher, B., Malone, K. M., Nguyen, G., Hall, M. B. et al. Minos: variant adjudication and joint genotyping of cohorts of bacterial genomes. Genome Biology 23, 147 (2022).

[37] Fowler, P. W. & Westhead, J. gnomonicus. https://github.com/oxfordmmm/gnomonicus (2023).

[38] Merker, M., Kohl, T. A., Barilar, I., Andres, S., Fowler, P. W. et al. Phylogenetically informative mutations in genes implicated in antibiotic resistance in Mycobacterium tuberculosis complex. Genome Medicine 12, 27 (2020).

[39] Fowler, P. W. & Westhead, J. piezo v0.8.1. https://github.com/oxfordmmm/piezo (2024).

[40] Pedregosa, F., Varoquaux, G., Gramfort, A., Michel, V., Thirion, B. et al. Scikit-learn: Machine learning in Python. Journal of Machine Learning Research 12, 2825–2830 (2011).

[41] Adlard, D. & Fowler, P. W. catomatic (2024). DOI: 10.5281/zenodo.14917986.

[42] Adlard, D. & Fowler, P. W. https://github.com/fowler-lab/tb-bdq-cat (2025).

[43] Vargas, R., Freschi, L., Marin, M., Epperson, L. E., Smith, M. et al. In-host population dynamics of Mycobacterium tuberculosis complex during active disease. eLife 10 (2021).

[44] MacLean, E. L.-H., Miotto, P., Angulo, L. G., Chiacchiaretta, M., Walker, T. M. et al. Updating the WHO target product profile for next-generation Mycobacterium tuberculosis drug susceptibility testing at peripheral centres. PLOS Global Public Health 3, e0001754 (2023).

[45] Snobre, J., Villellas, M. C., Coeck, N., Mulders, W., Tzfadia, O. et al. Bedaquiline- and clofazimine-selected Mycobacterium tuberculosis mutants: further insights on resistance driven largely by rv0678. Scientific Reports 13 (2023).

[46] Snobre, J., Meehan, C. J., Mulders, W., Rigouts, L., Buyl, R. et al. Frameshift Mutations in rv0678 Preserve Bedaquiline Susceptibility in Mycobacterium tuberculosis by Maintaining Protein Integrity (2024).

