## Supplemental Information for "An improved catalogue for whole-genome sequencing prediction of bedaquiline resistance in *M. tuberculosis* using a reproducible algorithmic approach"

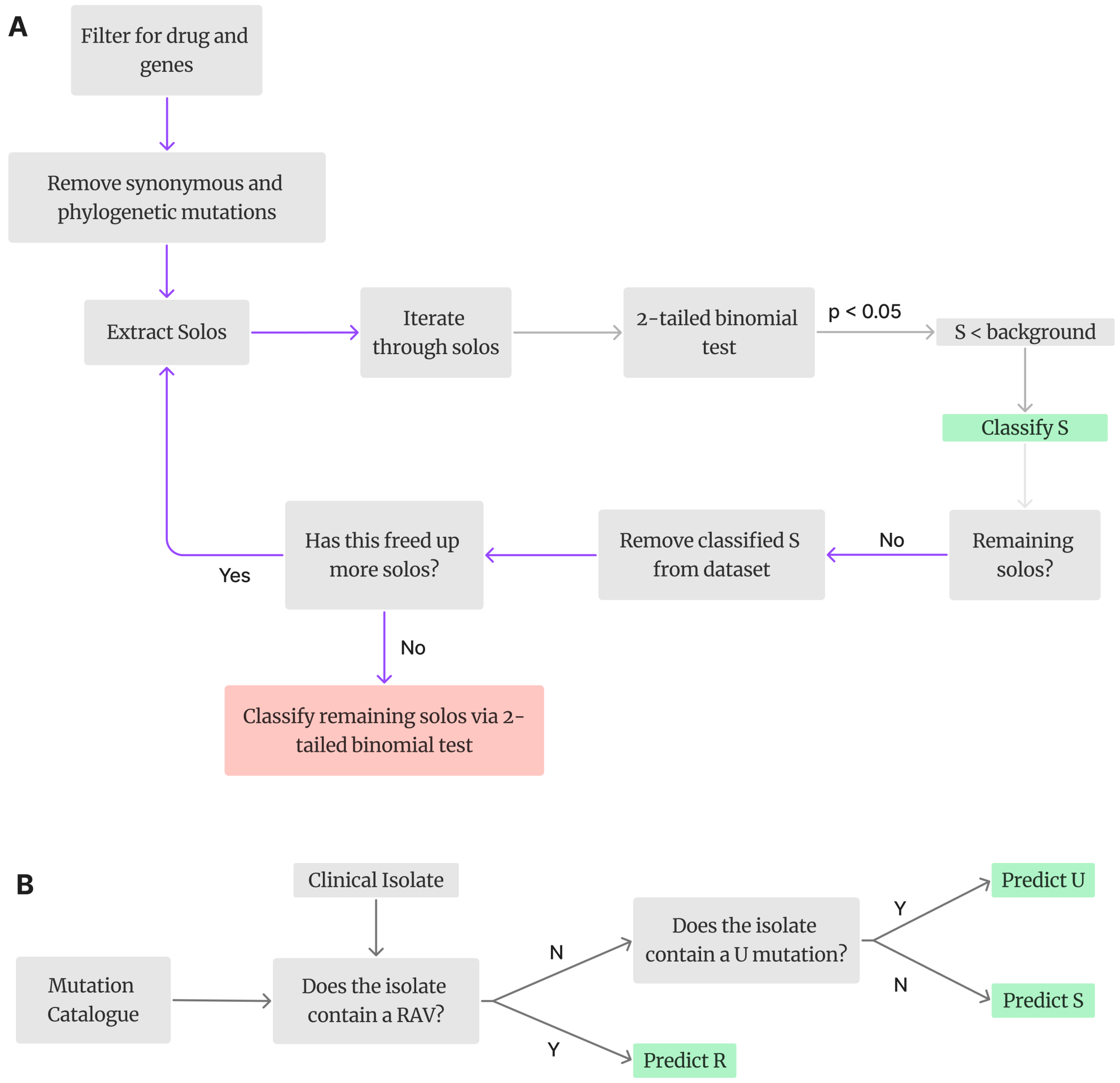


Figure S1: Decision workflow used to classify variants (A) and to make sample-level predictions using the *ternary* logical system (B). A. Mutations that exist in isolation across the candidate genes ("solos") are extracted, and a two-tailed binomial test determines whether the proportion of those alleles that are resistant is significantly different to a defined background rate of 10%. If the answer is yes in the direction of 0% resistance, then the mutation is classified as susceptible and removed from the dataset to proffer additional solo mutations. Once all susceptible mutations have been catalogued, the loop stops and all remaining variants are classified; if there are significantly more resistant alleles than the background, the mutation is classified R. If there is no significant difference, the mutation is classified U. B. If a sample contains a mutation catalogued as resistant, that sample is predicted as resistant to BDQ. If a sample contains an uncatalogued mutation or a mutation classified 'U', then it is predicted 'U'. Samples that contain no mutations in candidate genes, or mutations only catalogued 'S', are predicted to be susceptible.

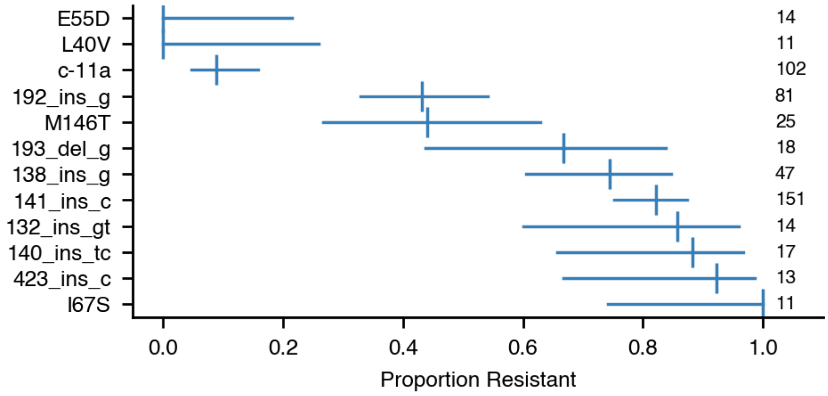


Figure S2: Proportion of resistance in samples that contain a high frequency mutation ( $>10$) in *Rv0678* as the only mutation across candidate genes, including minor variants, with confidence intervals, after removal of *mmpL5*, *mmpS5*, phylogenetic and synonymous mutations. By only considering mutations that exist in isolation in those samples, we constrain phenotypes to those only influenced by that mutation, which assumes we have indeed considered all resistance genes. mmpL5 and mmpS5 were removed as their mutations (of which there are many) are uniformly susceptible, and we thus logically assume they do not contribute to resistance. These proportions were extracted from the optimal catalogue referenced in the text (Table 3), in which binarised classification evidence is explicitly recorded.The heterogeneity observed, particularly for the indels, can be explained by examining their respective MIC distributions from which the labels were classified (Fig. S7). Were the background raised, such heterogeneity could pose a mis-classification risk.

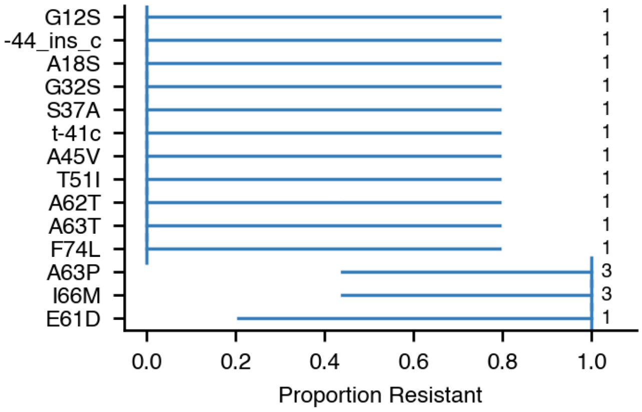


Figure S3: Proportion of resistance in samples that contain a mutation in *atpE* as the only mutation across candidate genes, with confidence intervals, after removal of *mmpL5*, *mmpS5*, phylogenetic and synonymous mutations, ordered by proportion of resistance. The same logic as Fig. S2 applies, without a minimum count threshold.

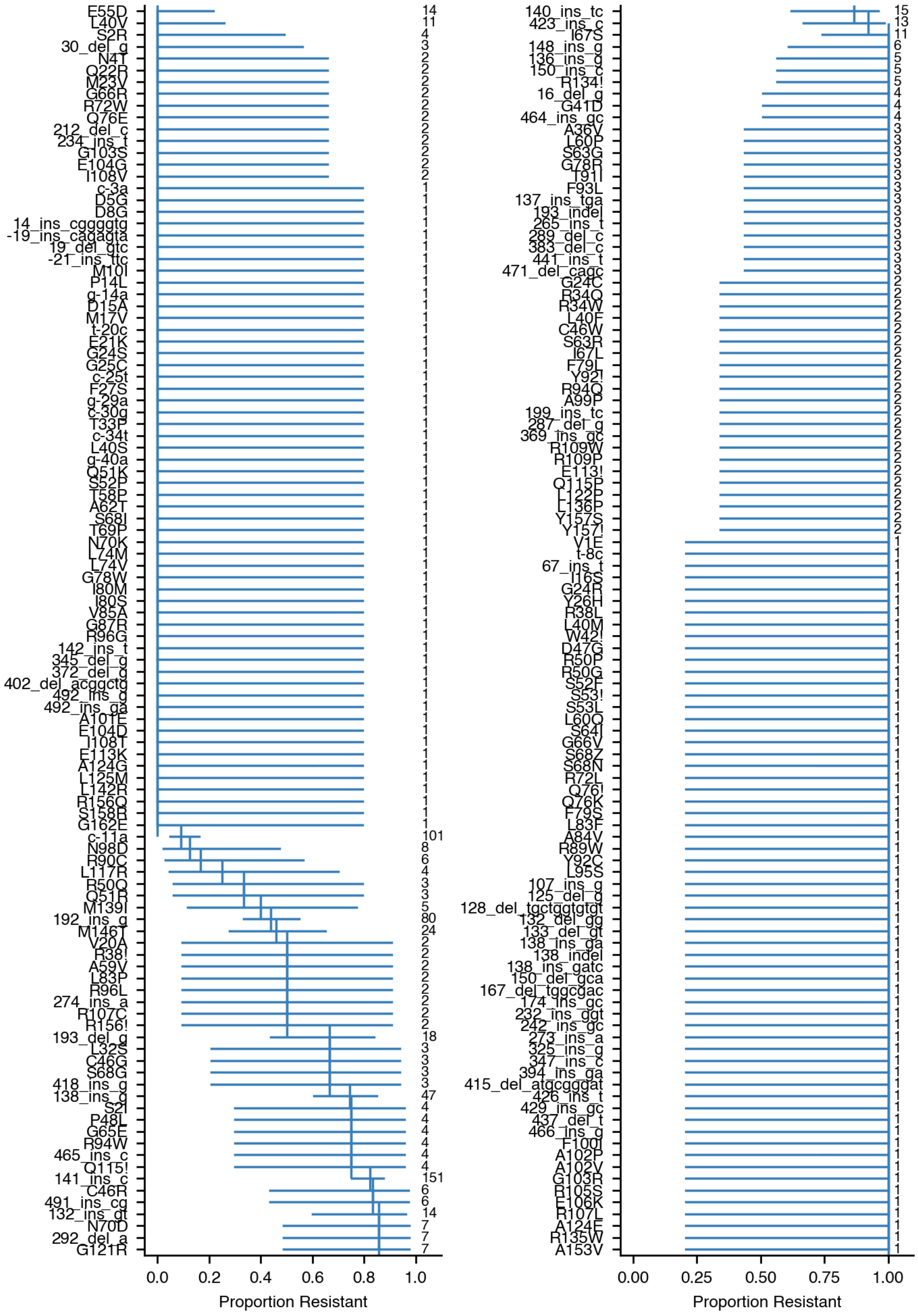


Figure S4: Proportion of resistance in samples that contain a mutation in *Rv0678* as the only mutation across candidate genes, with confidence intervals, after removal of *mmpL5*, *mmpS5*, phylogenetic and synonymous mutations, ordered by proportion of resistance. The same logic as Fig. S3 applies.

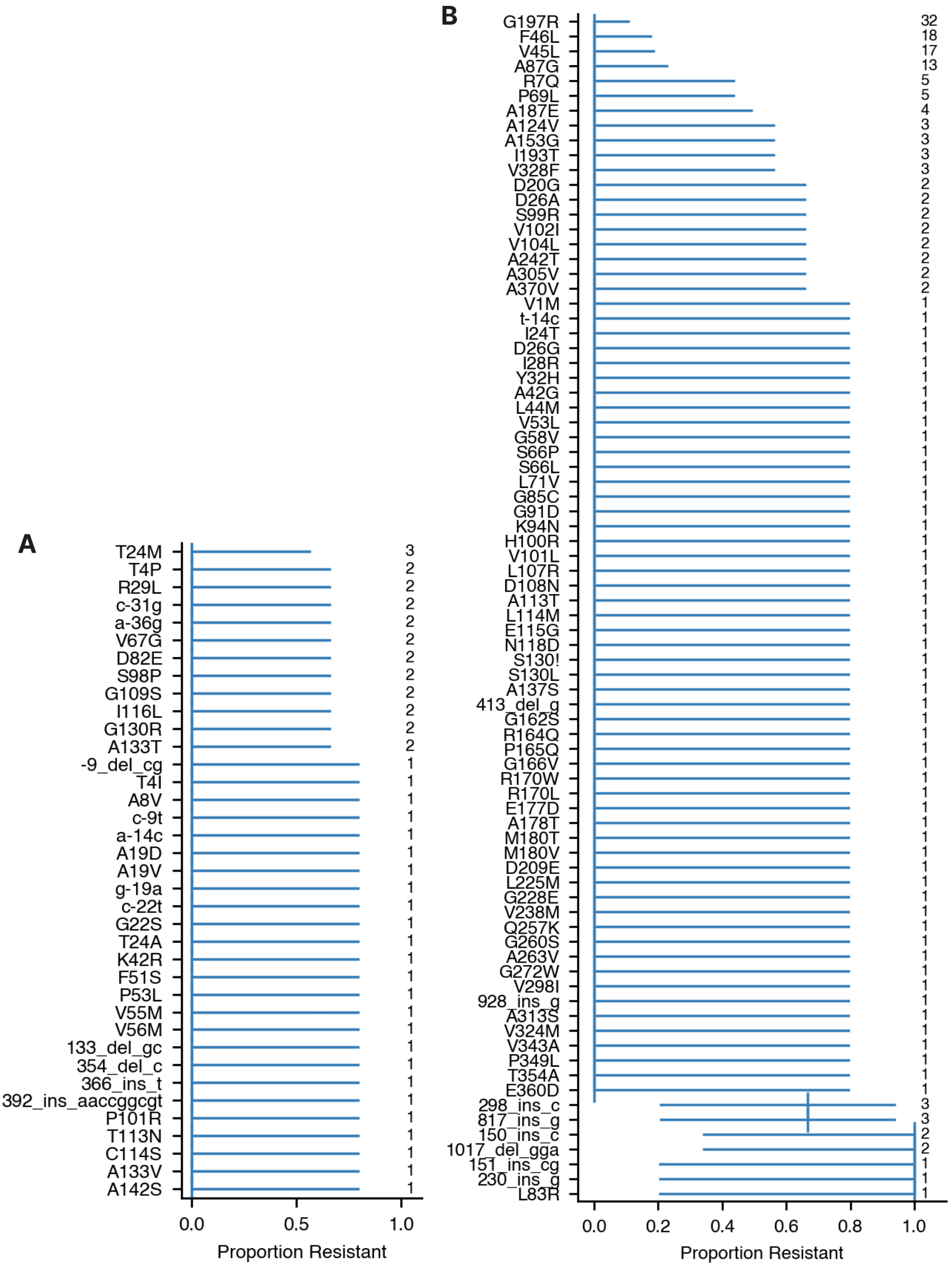


Figure S5: A.Proportion of resistance in samples that contain a mutation in *mmpS5* as the only mutation across candidate genes, after removal phylogenetic and synonymous mutations, ordered by proportion. B. Proportion of resistance in samples that contain a mutation in *pepQ* , after removal of *mmpL5*, *mmpS5*, phylogenetic and synonymous mutations.The same logic as Fig. S3 applies.

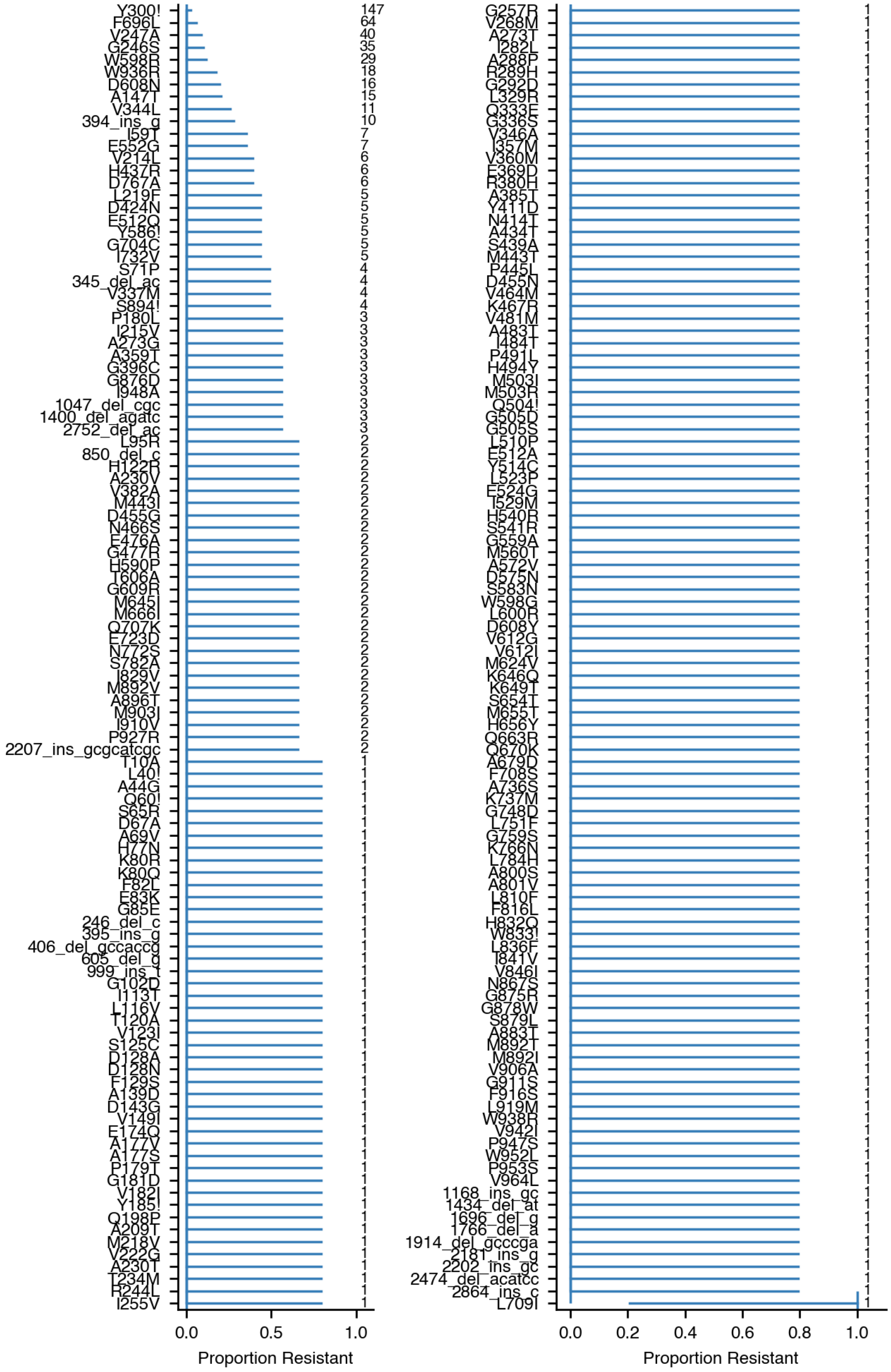


Figure S6: Proportion of resistance in samples that contain a mutation in *mmpL5* as the only mutation across candidate genes, with confidence intervals, after removal of phylogenetic and synonymous mutations, ordered by proportion of resistance. The same logic as Fig. S3 applies.

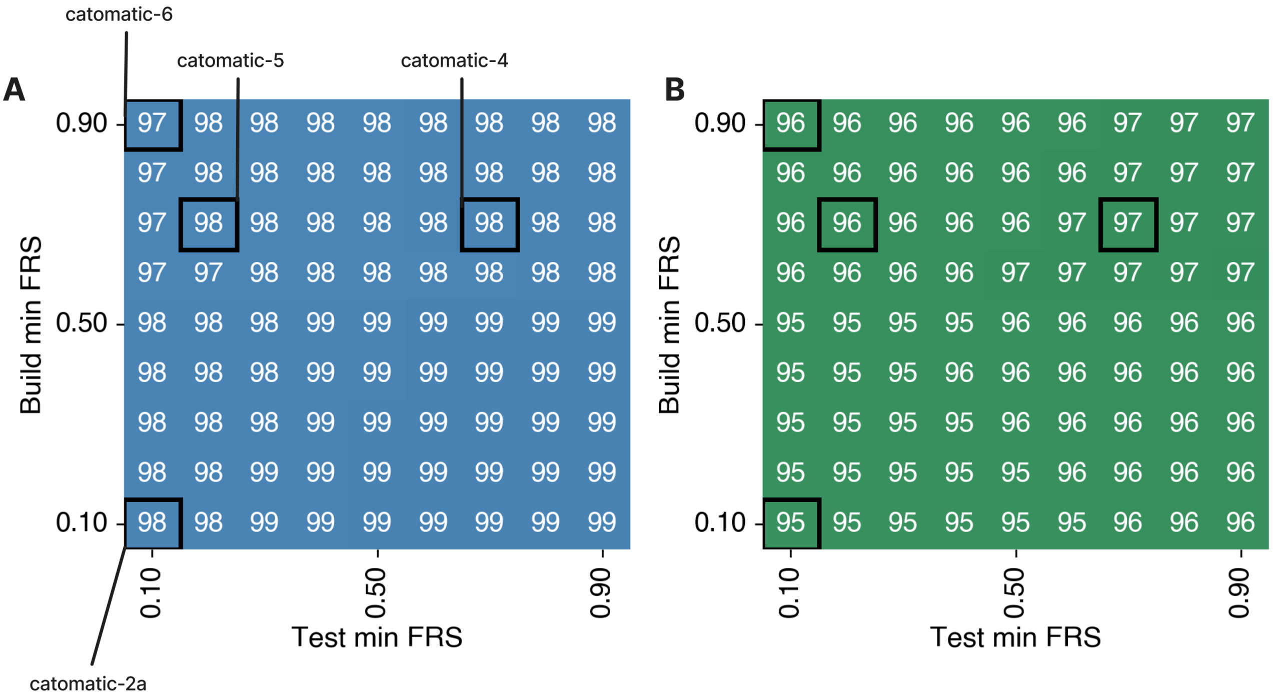


Figure S7: Specificity (A) and Definitive Prediction Rate (B) of the catomatic catalogue when evaluated *and constructed* as a function of minimum FRS at 0.10 increments, using the ternary prediction logic. Black boxes represent the 0.10 increments in which `catomatic-2a`, `catomatic-6`, `catomatic-5` and `catomatic-4`'s minimum FRS exist. These correspond to the sensitivity heatmap in Fig. 4F.

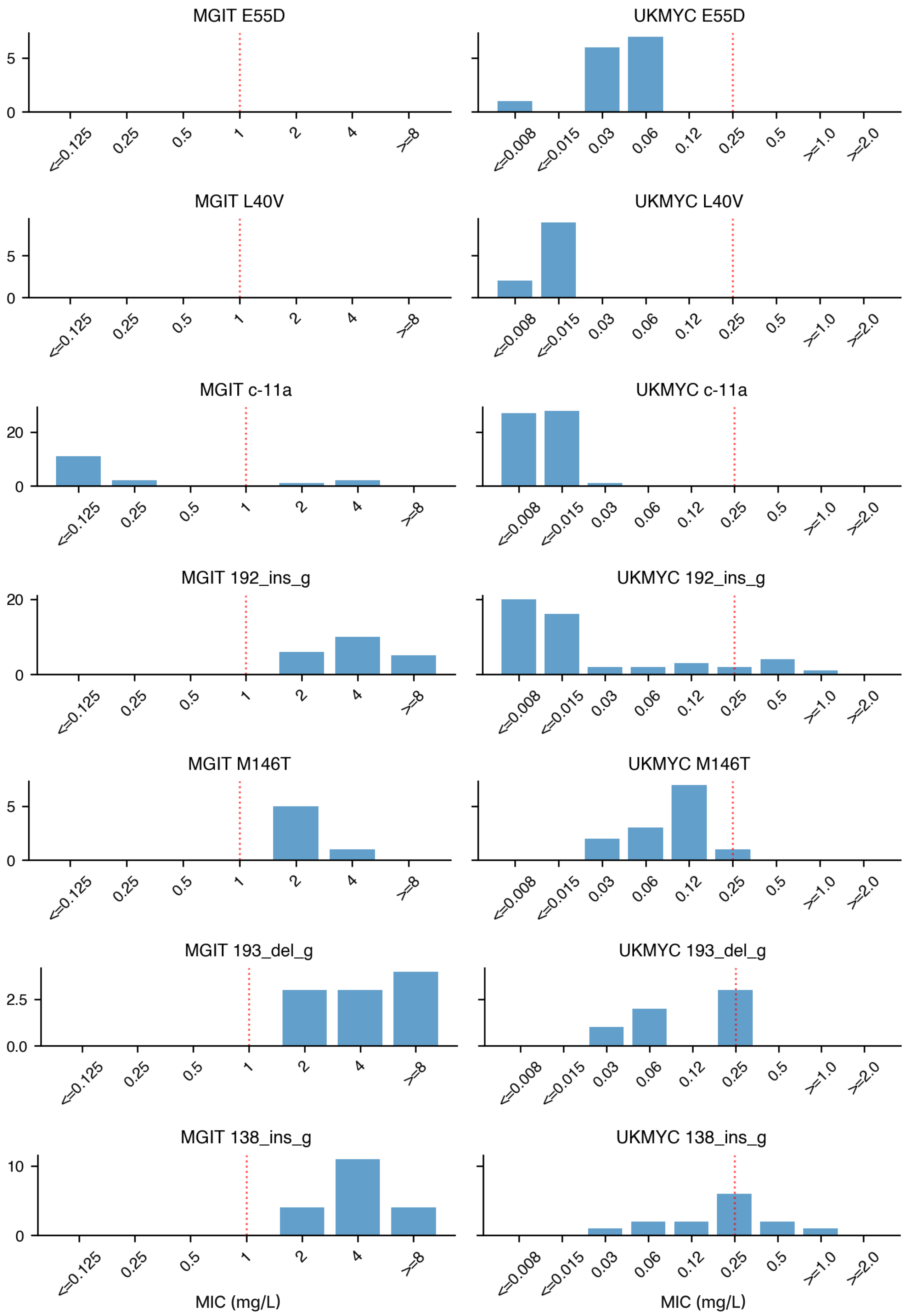
MGIT E55D
UKMYC E55D
MGIT L40V
UKMYC L40V
MGIT c-11a
UKMYC c-11a
MGIT 192_ins_g
UKMYC 192_ins_g
MGIT M146T
UKMYC M146T
MGIT 193_del_g
UKMYC 193_del_g
MGIT 138_ins_g
UKMYC 138_ins_g
<=0.125
0.25
0.5
1
2
4
>=8
<=0.008
<=0.015
0.03
0.06
0.12
0.25
0.5
>=1.0
>=2.0
MIC (mg/L)
MIC (mg/L)

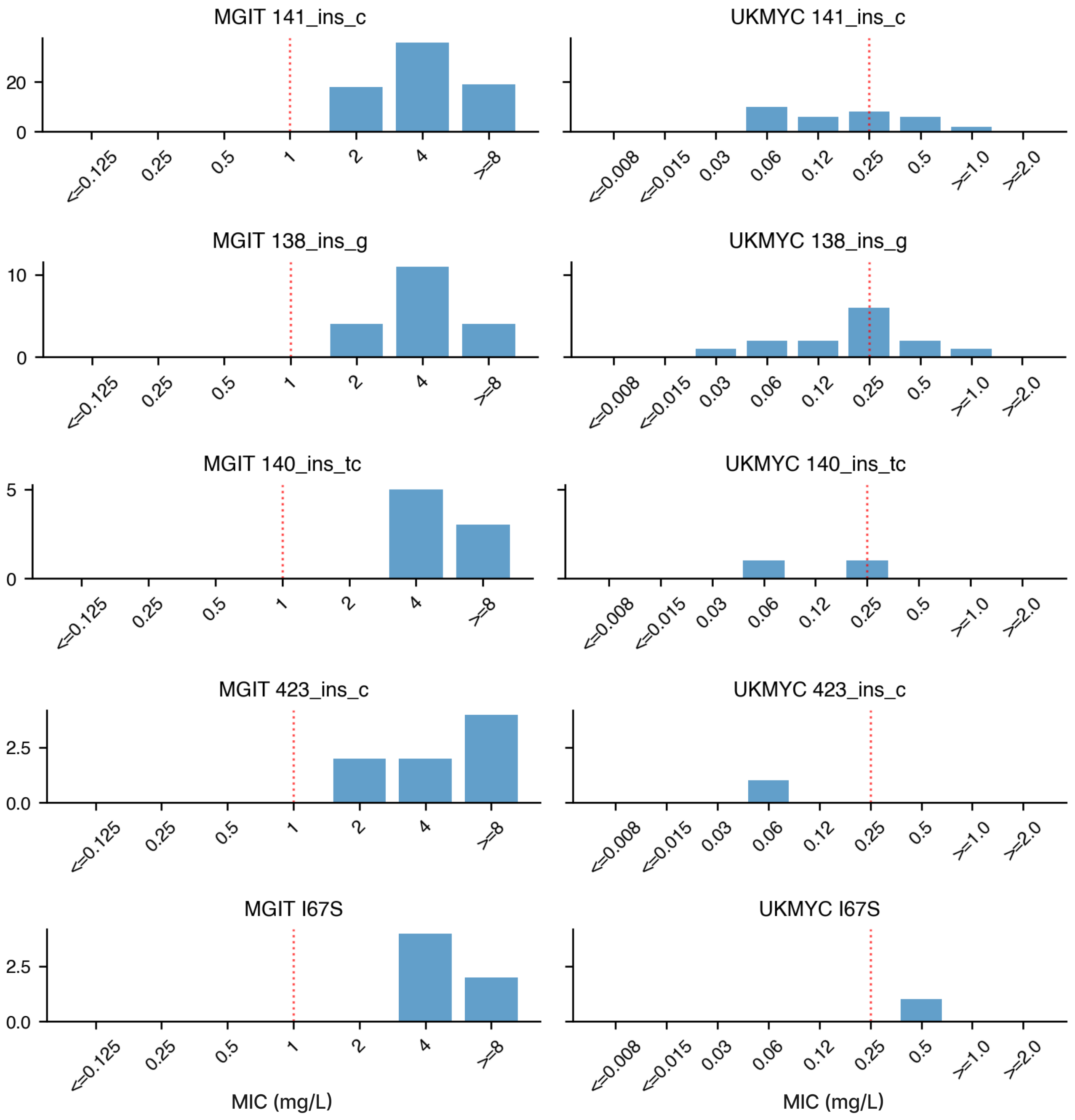


Figure S8: The minimum inhibitory concentration (MIC) distributions for each mutation in *Rv0678* observed more than 10 times in the training set. To avoid confounding only samples were this is the only nonsynoymous mutation in the resistance genes are considered. The ECOFF/ECV for each method is drawn as a red line; samples with an MIC above this threshold are classified as resistant (Fig. S2). Data collected using both the UKMYC5 and UKYMC6 plate designs have been aggregated. As mentioned in the Discussion, comparing the distributions is complex because the MGIT dataset is highly biased towards resistance and several of the samples with loss of functions mutations also have mutations in *mmpL5* which abrogate function leading to unusually low MICs. The differences could also be due to unaccounted for genetic effects, poorly calibrated ECOFF/ECVs, laboratory error or problems with one or both phenotyping methods. The relatively small increases in MIC for mutants observed using the UKMYC plates mirrors that seen for other non-essential genes, such as *embB*.

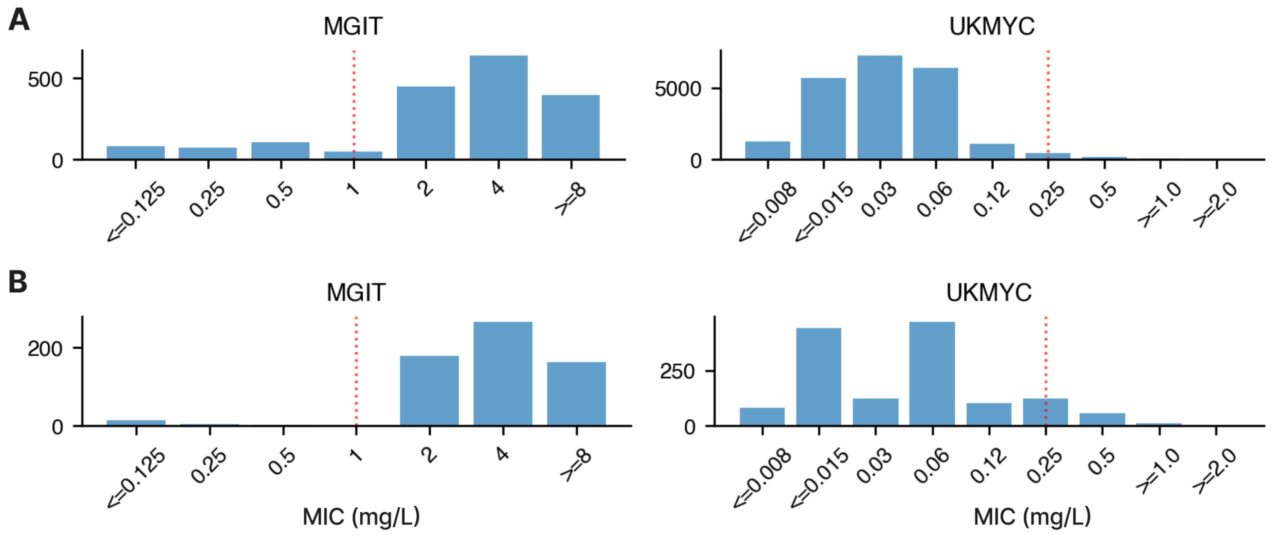


Figure S9: Minimum Inhibitory Concentration (MIC) distributions for all MGIT samples and UKMYC samples (A), and all non-WT samples with *mmpL5* and *mmpS5* mutations filtered out for MGIT and UKMYC samples (B). It is immediately clear basically all WT and low-MIC samples are provided by the UKMYC plates, and most high-MIC samples were phenotyped by MGIT. We accordingly need to combine the datasets to create a balanced training set, while recognising misaligned breakpoints (Fig. S8) could be a source of heterogeneity.
